# Nuclease-mediated depletion biases in ribosome footprint profiling libraries

**DOI:** 10.1101/2020.03.30.017061

**Authors:** Boris Zinshteyn, Jamie R Wangen, Boyang Hua, Rachel Green

**Affiliations:** Howard Hughes Medical Institute (HHMI); Department of Molecular Biology and Genetics, Johns Hopkins University School of Medicine, Baltimore, MD 21205

## Abstract

Ribosome footprint profiling is a high throughput sequencing based technique that provides detailed and global views of translation in living cells. An essential part of this technology is removal of unwanted, normally very abundant, ribosomal RNA sequences that dominate libraries and increase sequencing costs. The most effective commercial solution (Ribo-Zero) has been discontinued and a number of new, experimentally distinct commercial applications have emerged on the market. Here we evaluated several commercially available alternatives designed for RNA-seq of human samples and find them unsuitable for ribosome footprint profiling. We instead recommend the use of custom-designed biotinylated oligos, which were widely used in early ribosome profiling studies. Importantly, we warn that depletion solutions based on targeted nuclease cleavage significantly perturb the high-resolution information that can be derived from the data, and thus do not recommend their use for any applications that require precise determination of the ends of RNA fragments.

## Introduction

Ribosome footprint profiling (Ingolia et al. 2009; McGlincy and Ingolia 2017; Ingolia et al. 2019) provides nucleotide-precision snapshots of ribosome positions transcriptome-wide. This technique has been used by many groups to study a wide variety of biological problems, ranging from the mechanisms of translation regulation and mRNA quality control to questions about viral infection, stem cell differentiation, and mouse neurobiology. The technique works by sequencing the short (~30 nt) ribosome-protected mRNA fragments (footprints) that result from nuclease digestion of a cellular lysate (Wolin and Walter 1988; Kozak and Shatkin 1976; Steitz 1969). This nuclease digestion process inevitably causes widespread nicks in the ribosomal RNA (rRNA), producing an abundance of short rRNA fragments that copurify with intact ribosomes and comigrate with the ribosome footprints on polyacrylamide gels. Thus, the resulting sequencing datasets largely consist of rRNA sequences (Ingolia et al. 2009), limiting the sequencing depth of useful ribosome footprints.

Some early studies were able to mitigate contamination with rRNA fragments by gel purifying a very tight distribution of ~30 nt RNA fragments (the expected size of a ribosome footprint) to avoid prominent rRNA fragments (Guo et al. 2010), an approach which will not work for all species and samples. Cutting a narrow range of fragments also excludes other lengths of ribosome footprints, which report on the often substantial populations of ribosomes lacking A-site tRNAs (Lareau et al. 2014; Wu et al. 2019), trapped on truncated mRNAs (Guydosh and Green 2014; Arribere and Fire 2018; D’Orazio et al. 2019), or engaged in collisions (Guydosh and Green 2014).

Later studies used subtractive hybridization with biotinylated oligos to deplete the most abundant rRNA fragments (Brar et al. 2012; Ingolia et al. 2012), but variation in sample and library preparation can lead to differences in the number and identity of the most abundant contaminants, often requiring design of new probes with each modification of protocol. Commercial rRNA depletion solutions for RNA-seq are generally designed to work on intact or lightly fragmented rRNA, whose fragments are present in equimolar ratios. These assumptions do not apply to the rRNA fragments in ribosome profiling libraries, which are hugely biased by nuclease digestion and size selection. The standard rRNA depletion reagent for ribosome profiling became Ribo-Zero, which consisted of internally biotinylated and fragmented RNA antisense to the entire rRNA (Sooknanan 2009) and was able to deplete a broad range of fragments from ribosome profiling libraries, albeit with varying success (McGlincy and Ingolia 2017).

The recent discontinuation of Ribo-Zero led us to test several alternative commercial solutions designed for rRNA depletion of human samples. We found that the commercial products that we tested were at best only modestly effective at depleting rRNA from ribosome profiling libraries. More importantly, we found that methods utilizing targeted nucleolytic degradation of rRNA, such as by RNaseH, caused off-target trimming and degradation of ribosome footprints. Broadly speaking, these treatments reduce the fraction of mappable sequences, perturb global gene expression measurements and blur positional information.

## Results

In order to identify an alternative rRNA depletion option for ribosome footprint profiling we tested a small number of commercial depletion technologies designed for RNA-seq of human samples, alongside the remainder of our existing Ribo-Zero stock (hereafter referred to as legacy Ribo-Zero). In brief, our protocol (Wu et al. 2019; McGlincy and Ingolia 2017) consists of RNaseI digestion of cellular lysates, followed by pelleting ribosomes over a sucrose cushion and isolation of fragments by PAGE. To prepare them for sequencing (Figure 1A), these fragments are then ligated to a 3’ pre-adenylated DNA linker, depleted of rRNA fragments, reverse transcribed, circularized, and PCR amplified for Illumina sequencing. For this comparison, we prepared ribosome footprints from the same HEK293T cell lysate in duplicate, cutting broadly to isolate 15 to 40 nt fragments, which include the standard 30nt ribosome footprints, as well as the ~15-18 and 21 nt fragments that report on mRNA cleavage (Guydosh and Green 2014) and unoccupied A sites (Wu et al. 2019), respectively. We split these each into 4 identical groups after 3’ linker ligation (Figure 1A), to be depleted with Ribo-Zero (Illumina), RiboCop (Lexogen), NEBNext (NEB), or left undepleted. RiboCop is an affinity depletion method that uses complementary biotinylated DNA oligos targeting the rRNA. The NEBNext rRNA depletion kit uses antisense DNA oligos that tile the entire rRNA to target RNaseH-mediated degradation of rRNA. As described above, Ribo-Zero consists of biotinylated rRNA fragments complementary to entire rRNAs (Sooknanan 2009).

**Figure 1:**
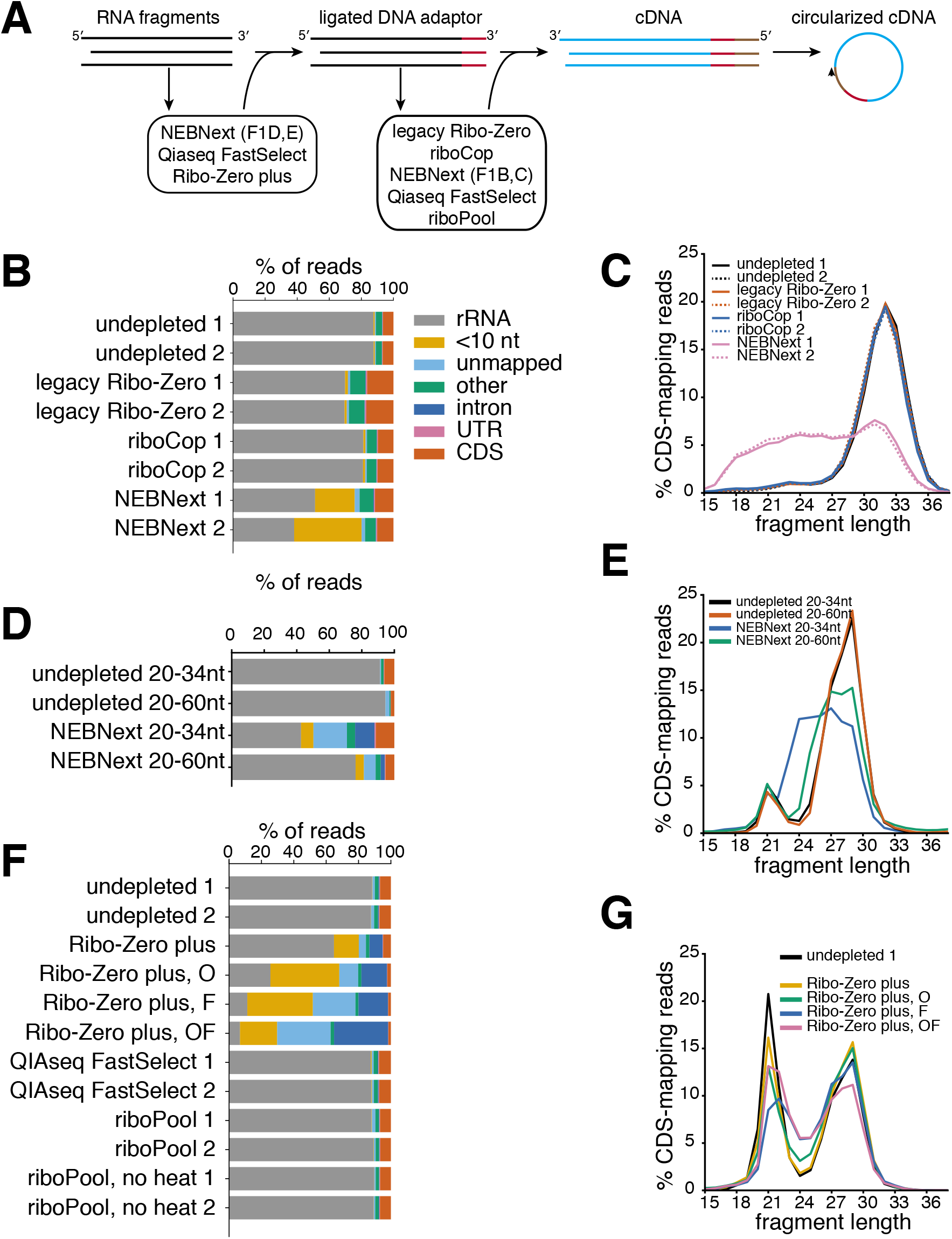
rRNA depletion and fragment length perturbation by commercial depletion kits. A) Simplified flowchart of library preparation for ribosome profiling, with steps at which various rRNA depletion strategies were performed indicated. Arrowhead on circularized cDNA indicates footprint 5’ end. B) Fractions of reads that are too short, unmappable, or mapping to various genomic regions for different rRNA depletion methods in HEK293T cells. C) Length distribution of ribosome footprints mapping to coding regions. D, E) Same as B and C for 2 different ranges of fragment size selection, depleted with the NEB kit in K562 cells. F, G) Same as B and C, for various depletions strategies in K562 cells. In panel G, only Ribo-Zero plus datasets are shown, as others did not have changes in length distribution.

After sequencing the resulting libraries, we found that the undepleted libraries contained 6.6±0.13% of reads mapping to coding regions (CDS) (Figure 1B). Legacy Ribo-Zero increased this to 16.6±0.33%, while RiboCop and NEBNext yielded 9.78±0.12% and 10.79±0.95%, respectively. While all of these approaches increased the number off useable reads, we noticed that the NEBNext kit produced a large fraction of short and unmappable fragments (“<10nt” and “unmapped” in Figure 1B). To investigate this further, we looked at the fragment size distribution in these libraries (Figure 1C). Ribosome footprint profiling in human cells normally produces fragments of ~30 nt, though the length distribution can be affected by RNaseI digestion conditions such as salt concentration (Ingolia et al. 2012). The undepleted libraries from this experiment have a modal fragment length of 32, with the majority of fragments between 28 and 36 nt in length. The libraries depleted by RiboCop and legacy Ribo-Zero have indistinguishable size distributions; importantly, however, the RNaseH-mediated depletion by the NEBNext protocol resulted in a broad distribution of fragment sizes spanning nearly the entire range of the 15 to 40 nt size selection. These data indicate that substantial off-target RNaseH cleavage has occurred. Since we performed RNA depletion immediately after linker ligation (Figure 1A), we surmised that this trimming was due to an abundance of leftover DNA linker, which contains 6 random nucleotides at its 5’ end that can anneal to our ribosome footprints and mediate RNAseH cleavage. To test this hypothesis, we repeated the profiling protocol, this time in K562 cells, with NEBNext depletion prior to linker ligation. We also tested the protocol with 2 different ranges of gel-purified fragments, 20-34 nt which corresponds to the footprint of a single ribosome, as well as 20-60 nt, which also includes the larger footprints of collided disomes (Guydosh and Green 2014), but which typically results in more rRNA contamination. The resulting depleted libraries contained substantially fewer short and unmappable reads than those produced with depletion post-ligation (compare Figure 1D to Figure 1B), suggesting that many of the short and unmappable fragments in our initial experiment were the result of widespread cleavage mediated by contaminating DNA linker. These datasets reveal a much tighter size distribution than post-ligation depletion, with peaks of ribosome-protected fragments 29 nt and 21 nt in length (Figure 1E), which correspond to ribosomes with occupied and unoccupied A sites, respectively (Wu et al. 2019). However, there was still noticeable trimming of the ribosome footprints, particularly the 29 nt fragments, which we presume is off-target activity from the DNA probes in the NEBNext kit.

We next tried three additional commercial rRNA depletion kits. These included a pre-release version of Ribo-Zero plus (Illumina) (a targeted depletion method similar to NEBNext that utilizes a proprietary nuclease), QIAseq FastSelect (Qiagen) (a pool of locked nucleic acids that block reverse transcription), and riboPOOL (siTOOLs biotech) (a cocktail of biotinylated oligos against the rRNA). We performed Ribo-Zero plus before linker ligation, and Qiaseq and RiboPOOL depletion immediately before reverse transcription (Figure 1A). We performed variations of Ribo-Zero plus depletion with supplementary oligos against abundant rRNA fragments (O), as well as supplemented with formamide (F) during probe annealing, to better denature highly structured rRNA fragments. For riboPOOL depletion, we also performed a variation where we omitted the final heating step. None of these methods proved effective at increasing the fraction of CDS-mapping fragments (Figure 1F). Although Ribo-Zero plus decreased the fraction of rRNA-mapping reads, it appears that CDS-mapping reads were degraded, and the library contained an abundance of unmappable or intronic reads. These effects are evident in the CDS-mapping footprint length distribution (Figure 1G), which shows a broadening of the 21 and 29 nt peaks and a visible increase in the fraction of 23-25 nt reads, which likely are degradation product of the longer 28nt fragments, that correlates with the extent of depletion. This fraction increases from 7% in the undepleted sample to 20% in the Ribo-Zero plus OF library. We interpret this result as indicating that off-target activity is a general feature of nuclease-mediated rRNA depletion.

Since the NEBNext kit was modestly successful at rRNA depletion, we tested if these depleted libraries were still suitable for making gene-level inferences of ribosome footprint density. The RPM (reads per million CDS-mapping reads) values for our undepleted or NEBNext depleted libraries were reproducible (Pearson r^2^ = 0.900 and 0.922) between pseudoreplicates (pseudo because we isolated different fragment sizes) despite their modest sequencing depth, but many transcript RPMs differed between the depleted and undepleted libraries (Pearson r^2^ = 0.788 and 0.780), with a small number of genes showing extreme deviations in RPM (Figure 2A). These differences indicate that even aggregate gene-level measurements are perturbed by RNaseH-mediated depletion. This perturbation was also seen in the Ribo-Zero plus depleted samples (Figure 2B), suggesting that it is a general feature of nuclease-mediated depletion.

**Figure 2:**
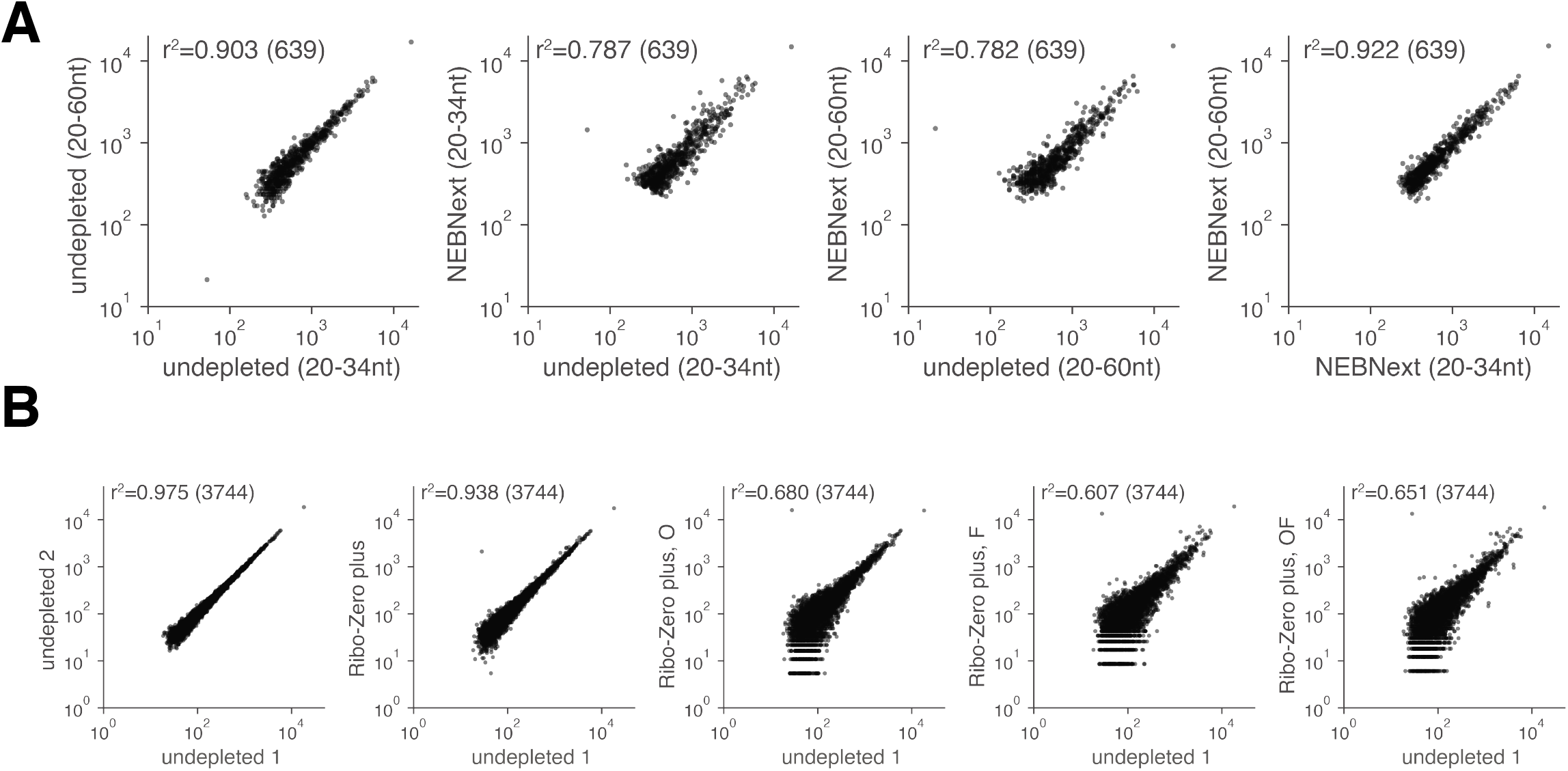
effect of nuclease-mediated rRNA depletion on gene-level expression measurements. Correlations between CDS-mapping RPMs (reads per million) for replicates, and for libraries depleted with A) NEBNext or B) Ribo-Zero plus. Pearson correlation of log-transformed values is indicated.

An important feature of ribosome profiling data is the nucleotide-resolution positional information, which provides information on ribosome movement at specific codons or groups of codons (Ingolia et al. 2009; Stadler and Fire 2011; Zinshteyn and Gilbert 2013; Nedialkova and Leidel 2015; Ingolia et al. 2011), as well as the A site status (occupied or empty) of the ribosome, which is inferred from the length of the footprint (Wu et al. 2019). These analyses require the identification of the A, P and E site locations within each footprint, as well as the length class (~21, 28, etc.) of that footprint. Site identification is generally accomplished by averaging reads of a given length relative to all start codons, with the knowledge that ribosomes initiate with the start codon in the P site. Both NEBNext and Ribo-Zero plus depletion blurred the accuracy of site assignment (Figure 3) as evidenced by reduced peaks and shallower troughs in the average gene plots.

**Figure 3:**
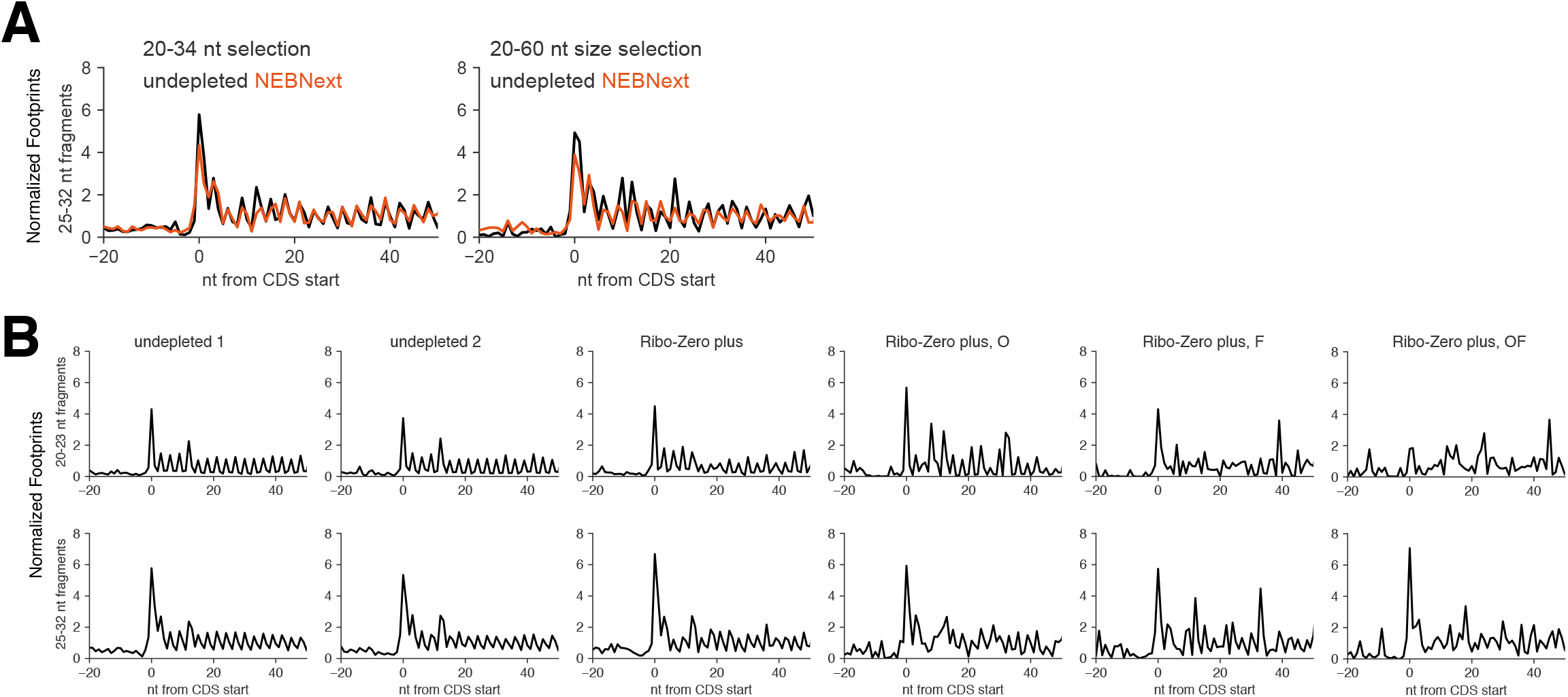
average gene profiles for depleted datasets. Average gene plots aligned at start codons for libraries depleted with A) NEBNext or B) Ribo-Zero plus. Reads have been offset to indicate ribosomal P sites. Reads are normalized by the read density of their respective coding region, such that all genes with sufficient read numbers contribute equally, regardless of expression level. The Ribo-Zero O, F and OF datasets have very low numbers of CDS-mapping reads, leading to low-quality average gene plots.

Ribosome profiling libraries with perturbed read lengths could still be useable for positional analysis if the read trimming happens in a defined way. To see how the alterations of read lengths affect the positional resolution of ribosome profiling, we summed the total number of footprint 5’ ends for each read length relative to the start codons of all coding regions (Figures S1 and S2). Ribosome profiling libraries generally have a peak of footprint 5’ ends mapping approximately 12 nt upstream of start codons. In the 5’-aligned footprint heatmaps in Figures S1 and S2, diagonal lines from the start codon peak are characteristic of footprints from initiating ribosomes that are shortened by nibbling at the 5’ end, while vertical lines indicate variation in the 3’ end, which maintains the position of the 5’ end while shortening the footprint. The presence of both vertical and diagonal lines emanating from the start codon peaks of the NEBNext-depleted libraries (Figures S1B and S2) indicates that the reads are trimmed from both ends. Since it is impossible to determine from which end a particular read is truncated, this trimming is unlikely to be computationally correctable.

In a final experiment, we decided to revisit previously used methods for rRNA depletion based on antisense biotinylated oligos (Ingolia et al. 2012). Based on the sequencing data from our undepleted K562 samples, we designed a set of 6 biotinylated oligos complementary to 37% of rRNA fragments (Table S1). We used these oligos to subtract rRNA fragments from adaptor-ligated footprints generated from HEK293T cells. This step increased the fraction of CDS-mapping reads in our library from 5.7% to 8.7%, with little or no perturbation of fragment lengths, positions, or gene-level fragment counts (Figure 4). These promising results suggest that further efforts for ribosome profiling rRNA depletion should be aimed at optimizing custom oligo-based methods.

**Figure 4:**
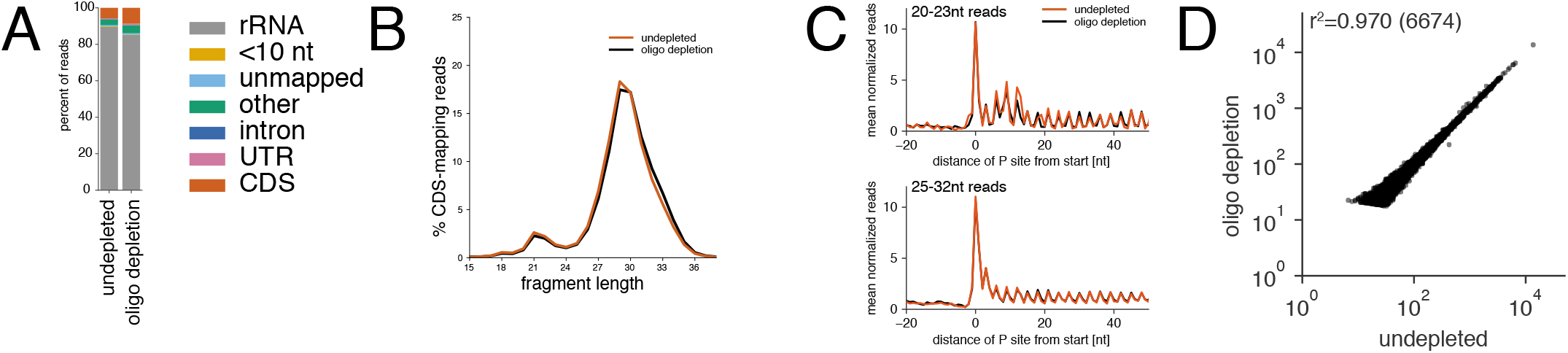
rRNA depletion with targeted biotinylated oligos. A) Fractions of reads that are too short, unmappable, or mapping to various genomic regions for antisense oligo rRNA depletion in HEK293FT cells. B) Length distribution of ribosome footprints mapping to coding regions. C) Average gene plots aligned at start codons for undepleted libraries or oligo depleted libraries, broken up by read length. D) Correlations between CDS-mapping RPMs for undepleted libraries or oligo depleted libraries.

## Discussion

In summary, we have tried several rRNA depletion methods for ribosome profiling and find all of the commercially available options lacking compared to legacy Ribo-Zero. It is possible that with some optimization, some of these kits (most notably RiboCop) could be successfully used for human samples, but we strongly caution against the use of any methods that uses nuclease cleavage, as they blur the detailed mechanistic information that can be gained from the lengths of ribosome footprints (Wu et al. 2019; Guydosh and Green 2014). Moreover, most of these reagents are only available for use in a limited number of species. The best way forward for rRNA depletion in ribosome profiling may be the previously used custom oligo depletion methods (Ingolia et al. 2012), perhaps combined with biophysical separation methods, such as separating the intact ribosomes from the ribosome footprints. In yeast library preparations, we have had reasonable success using previously-published oligo depletion methods (Guydosh and Green 2014), consistently achieving 40% CDS-mapping reads with libraries cut from 20-34 nt (data not shown). However, our results have been less effective for human samples (Figure 4), indicating that further species-specific optimization in either depletion protocols or depletion oligo design is required. Laub and coworkers recently developed an effective algorithm for designing depletion oligos targeting multiple bacterial species for RNA-seq (Culviner et al. 2020), but it remains to be seen whether the same method can be adapted to ribosome profiling, which has a much more biased distribution of rRNA fragments than RNA-seq. Other groups have had success using double-strand specific DNase, combined with a high-temperature re-annealing of the final DNA library that only leaves the most abundant sequences (the rRNA fragments) doublestranded (Chung et al. 2015), but this method has not to our knowledge been tested on a broad range of fragment lengths, nor with improved library preparation methods such as randomized unique molecular identifiers (McGlincy and Ingolia 2017). Recent reports indicate that optimization of RNase digestion conditions (Sharma et al. 2019) as well as judicious choice of RNase enzyme (Gerashchenko and Gladyshev 2017) can also reduce the rRNA content of ribosome profiling libraries, though many procedures using these different nucleases obscure the high resolution information on ribosome position and A site status. Finally, it is important to note that off-target RNaseH activity will not only affect the conclusions of ribosome footprint profiling experiments, but of any protocol that requires precise knowledge of the 5’ and 3’ ends of RNA fragments.

## Methods

### Cell culture Conditions

Mammalian cell lines were grown in cell culture incubators at 37°C in the presence of 5% CO_2_. HEK293T cells were cultured in in Dulbecco’s modified Eagle media (DMEM) +10% fetal bovine serum (FBS) (Gibco). K562 cells were grown in Roswell Park Memorial Institute (RPMI) 1640 media + 10% FBS.

### Cell lysis and ribosome footprint profiling

Our protocol is adapted from (McGlincy and Ingolia 2017). 10 cm dishes of adherent HEK293T cells were briefly washed with PBS and lysed by addition of 0.5 mL of mammalian footprint lysis buffer [20 mM Tris-Cl (pH8.0), 150 mM KCl, 5 mM MgCl2, 1 mM DTT, 1% Triton X-100, 2 units/ml Turbo DNase (Thermo Fisher), 0.1 mg/mL cycloheximide (Sigma-Aldrich)] and vigorous scraping. 5-10 million K562 cells were harvested by centrifugation at 500*g* for 2 minutes at 37°C, washed briefly in 1 mL 37°C PBS, and lysed by pipetting in 200 μL mammalian footprint lysis buffer. Lysates were incubated on ice for 10 minutes and then clarified by centrifugation at >15,000*g* for 10 minutes. 0.5 A260 units of lysate was digested with 750 units of RNase I (Ambion) in 300 μL lysis buffer at 25°C for 1 hour. Reactions were quenched with 200 U SUPERase-in (Thermo Fisher) and pelleted over a sucrose cushion for 1 hr at 100,000 RPM. Pellets were resuspended in TRIzol (Thermo Fisher) and RNA extracted with the Zymo Direct-zol RNA miniprep kit. Fragments were size selected on a 15% TBE-urea PAGE gel (BioRad), cutting between RNA markers of 15-40, 20-34, or 20-60, depending on the experiment. After gel elution, fragments are dephosphorylated, ligated to preadenylated 3’ linker oBZ407 (AppNNNNNNCACTCGGGCACCAAGGAC/3ddC/), reverse transcribed using protoscript II (NEB) and RT primer oBZ408 (/5Phos/RNNNAGATCGGAAGAGCGTCGTGTAGGGAAAGAGTGTAGATCTCGGTGGTCGC/iSP18/TTC AGACGTGTGCTCTTCCGATCTGTCCTTGGTGCCCGAGTG), circularized with Circ Ligase (Lucigen), and PCR amplified. Amplified libraries were quantified by bioanalyzer high-sensitivity DNA chip (Agilent), pooled and sequenced on an Illumina HiSeq 2500 with 50 or 100 nt reads.

### rRNA depletion methods

For Figure 1A and 1B, Legacy Ribo-zero (Illumina), RiboCop (Lexogen) and NEBNext (NEB) depletions were performed after linker ligation according to the manufacturer’s recommendations. For Ribo-Zero depletion, we omit the final heating step in the manufacturer protocol, as this has been suggested to improve depletion of small fragments (McGlincy and Ingolia 2017). For subsequent figures NEBNext and Ribo-Zero plus (Illumina) depletion were performed right after fragment size selection, while all other methods were performed after linker ligation. These methods were performed according to manufacturer recommendations, except that for some experiments with Ribo-Zero plus we included 45% formamide in the hybridization reaction (indicated as F), as well as supplementary oligos (indicated as O), provided by Illumina, designed against abundant rRNA contaminants. For custom oligo depletion, an oligo mix solution was made by diluting the 6 depletion oligos (Table S1) in 4X SSC buffer to a concentration of 2.5 μM for each oligo. 10 μl of the oligo mix solution was added to 10 μl of sample solution (ligation products precipitated and resuspended in nuclease-free water). The sample and oligo mixture was heated at 80 °C for 2 min in a thermomixer, which was then set to 25 °C to gradually cool the mixture and anneal oligos to the sample RNA. During cooling, 150 μl of MyOne Streptavidin C1 dynabeads (ThermoFisher) per depletion were washed thrice with 150 μl of 1X Binding/Washing buffer (5 mM Tris, pH 8.0, 1 M NaCl, 0.5 mM EDTA), and then resuspend beads in 30 μl 2X Binding/Washing buffer. The cooled sample was added to the prepared beads and incubated at 25 °C for 15 min with shaking at 500 rpm in a thermomixer. After incubation, beads were precipitated with a magnetic stand, and supernatant was removed and isopropanol precipitated to recover RNA.

### Data Analysis

Raw ribosome footprint reads were trimmed of the 4 random nt from the 5’ end with seqtk (https://github.com/lh3/seqtk), then trimmed of 3’ adaptor sequence (NNNNNNCACTCGGGCACCAAGGAC) using Skewer (Jiang et al. 2014). Reads longer than 10nt were aligned to PhiX-174, human rRNA and human non-coding RNA sequences using STAR (Dobin et al. 2013) with --outFilterMismatchNmax 2 --outSAMmultNmax 1. Unmapped reads were mapped to the hg38 human genome (GENCODE release 27), using STAR with parameters --alignSJDBoverhangMin 1 --alignIntronMax 1000000 --alignSJoverhangMin 3 --outFilterType BySJout --outFilterMultimapNmax 200, --outFilterScoreMinOverLread 0 --outFilterMatchNminOverLread 0 --outFilterMatchNmin 0, --outFilterMismatchNmax 3. To generate a reference transcriptome with a single transcript per gene for alignment, the GENCODE v27 gtf file was filtered for those transcripts with an APPRIS score (Rodriguez et al. 2013) between 1 and 4 (inclusive) and then the transcript with the longest CDS (and longest transcript if tied) was chosen. For multiply-mapping reads, the primary alignment was used for all analyses. For Figures 1A, 1C and 1E, reads were assigned the smallest feature that hey overlapped based on the comprehensive GENCODE v27 annotation gtf file. For RPM correlation plots, only genes with an average of 64 or more counts and a minimum of 1 RPM between all libraries in the same panel were included. The Python code for the analysis pipeline and for figure generation are available at https://github.com/borisz264/rRNA_depletion_2020. Sequencing data have been deposited in GEO with accession GSE147324.

## Acknowledgements

We thank David Mohr and the Johns Hopkins Genetic Resources Core Facility for sequencing assistance. We thank Scott Kuersten (Illumina) for providing Ribo-Zero plus material as well as advice on its use, and Sezen Meydan for suggestions on the use of Qiagen FastSelect. This work was funded by the National Institutes of Health [2R37GM059425-14 to R.G.; 5K99GM135450-02 to B.Z], Howard Hughes Medical Institute (HHMI) (R.G.) and the Cystic Fibrosis Foundation (GREEN16G0). B.Z. was an HHMI fellow of the Damon Runyon Cancer Research Foundation [DRG-2250-16] for a portion of this study.

---

**Supplementary Figure 1:**
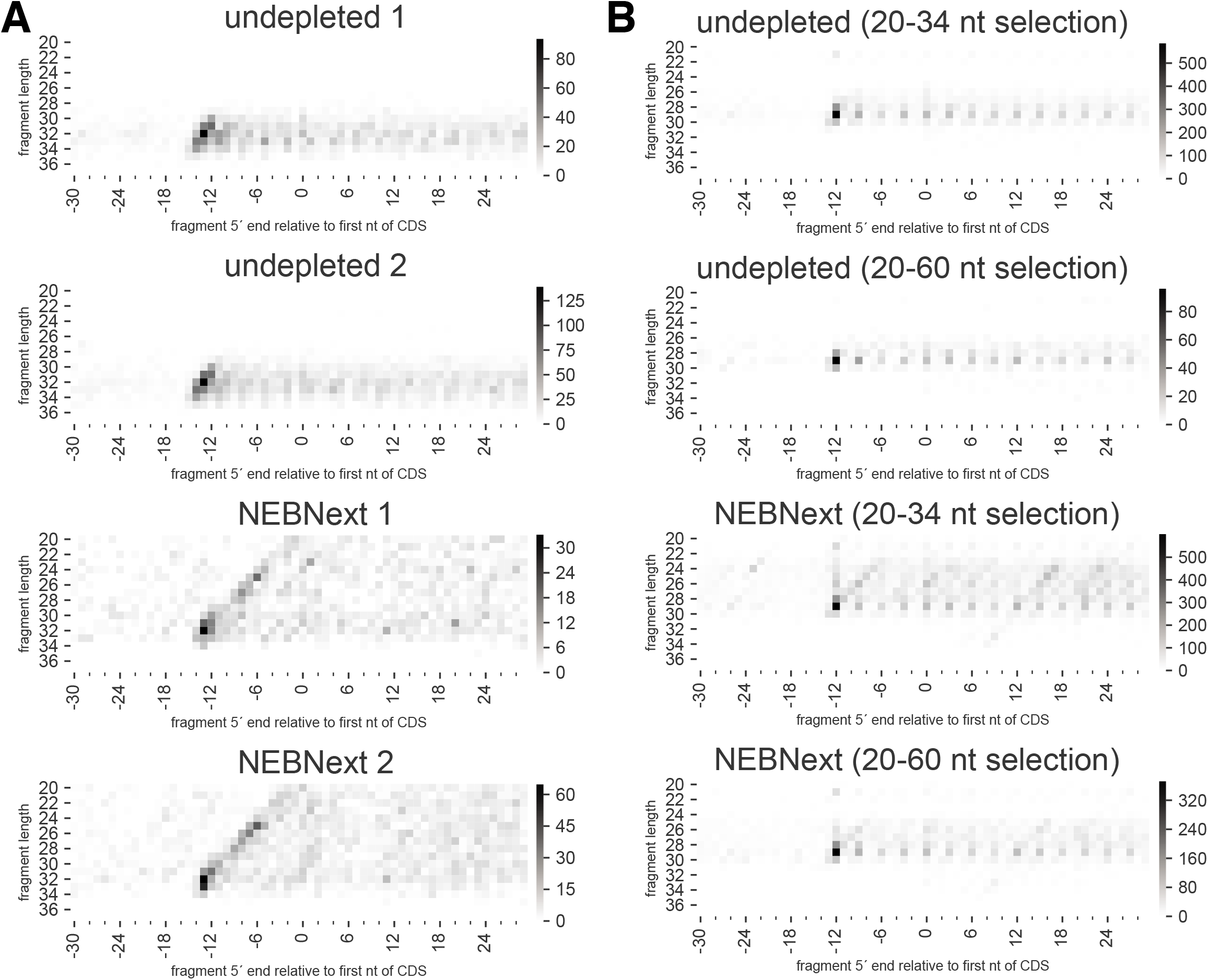
Fragment 5’ ends mapping to CDS starts by read length for NEBNext depletion. Heatmaps of number of ribosome footprint 5’ ends mapping relative to the first nucleotide of coding regions, subdivided by read length, for NEBNext depleted libraries from A) Figure 1A and B) Figure 1C.

**Supplementary Figure 2:**
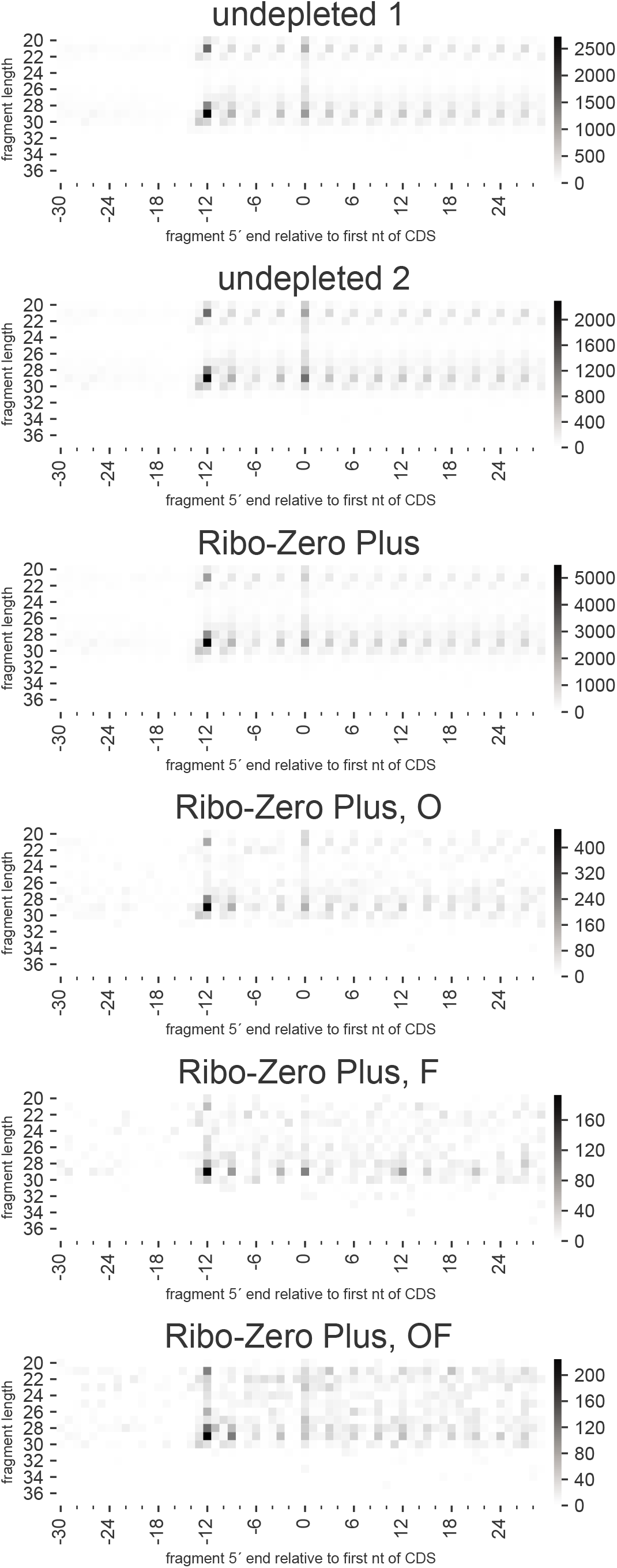
Fragment 5’ ends mapping to CDS starts by read length for Ribo-Zero plus depletion. Heatmaps of number of ribosome footprint 5’ ends mapping relative to the first nucleotide of coding regions, subdivided by read length, for Ribo-Zero Plus depleted libraries from Figure 1E.

